# Prospects of genomic prediction in the USDA Soybean Germplasm Collection: Historical data creates robust models for enhancing selection of accessions

**DOI:** 10.1101/055038

**Authors:** Diego Jarquin, James Specht, Aaron Lorenz

## Abstract

The identification and mobilization of useful genetic variation from germplasm banks for use in breeding programs is critical for future genetic gain and protection against crop pests. Plummeting costs of next-generation sequencing and genotyping is revolutionizing the way in which researchers and breeders interface with plant germplasm collections. An example of this is the high density genotyping of the entire USDA Soybean Germplasm Collection. We assessed the usefulness of 50K SNP data collected on 18,480 domesticated soybean (*G. max*) accessions and vast historical phenotypic data for developing genomic prediction models for protein, oil, and yield. Resulting genomic prediction models explained an appreciable amount of the variation in accession performance in independent validation trials, with correlations between predicted and observed reaching up to 0.92 for oil and protein and 0.79 for yield. The optimization of training set design was explored using a series of cross-validation schemes. It was found that the target population and environment need to be well represented in the training set. Secondly, genomic prediction training sets appear to be robust to the presence of data from diverse geographical locations and genetic clusters. This finding, however, depends on the influence of shattering and lodging, and may be specific to soybean with its presence of maturity groups. The distribution of 7,608 non-phenotyped accessions was examined through the application of genomic prediction models. The distribution of predictions of phenotyped accessions was representative of the distribution of predictions for non-phenotyped accessions, with no non-phenotyped accessions being predicted to fall far outside the range of predictions of phenotyped accessions.

## 1. Introduction

The foundation of plant breeding is genetic diversity yet the success of modern scientific plant breeding is leading to an erosion of the very genetic diversity it relies upon as farmers discard landraces in favor of genetically improved and uniform cultivars derived from a limited ancestral base. This genetic erosion increases vulnerability to agricultural insect and disease epidemics, as well as diminishes gains from breeding and selection. Germplasm collections serve as an important source of variation for germplasm enhancement; that variation sustains long-term genetic gain in breeding programs. A stunning number of accessions −7.4 million -- is being maintained *ex situ* by plant germplasm collections worldwide, also referred to as gene banks (FAO[Food and Agriculture Organization] 2010). The largest number of accessions belongs to wheat with approximately 856,000 accessions held, followed by rice with nearly 774,000 accessions (FAO[Food and Agriculture Organization] 2010). The USDA National Plant Germplasm System (NPGS) alone holds more than 571,207 accessions for 14,965 species as of June 2015, ranging from 53,525 accessions for rice to 165 accession for quinoa(http://www.ars-grin.gov/npgs/stats/summary.html).

The identification and mobilization of useful genetic variation from germplasm banks for use in breeding programs is clearly a necessity not only for sustaining current rates, but also for increasing future rates of crop genetic improvement (Sehgal *et al.* 2015). Nevertheless, there is evidence that these collections are woefully underutilized. In 2004, Carter and coworkers estimated that among approximately 45,000 unique soybean accessions maintained in germplasm collections worldwide, only 1,000 have been used in applied breeding programs (Carter *et al.* 2004). Beneficial alleles for traits like yield have been mined from exotic and wild germplasm (Tanksley *et al.* 1996; Fox *et al.* 2015), and breeders accept that landraces and exotic germplasm likely contain alleles that could enhance their germplasm, even for intensely selected traits, such as yield. However, efficiently mining such large germplasm collections with little knowledge on accession breeding values and the distribution of favorable alleles for complex traits like yield is a huge challenge, yet selecting exotic parents for yield improvement is just as critical as selecting elite parents.

Plummeting costs of next-generation sequencing (NGS) is revolutionizing the way in which researchers and breeders interface with plant germplasm collections. It is possible that all accessions held worldwide will be densely genotyped using NGS technologies. Some present examples of wide-scale genotypic characterization of the germplasm collections include the genotyping by sequencing of the CIMMYT maize collection (Hearne *et al.* 2015) and the sequencing of 3,000 rice genomes (Li *et al.* 2014). This information will greatly benefit the selection of accessions for breeding and genetics research. Using genomic data, accessions could be selected which contain specific alleles of desired effect (McCouch *et al.* 2012), or all accessions representing all allelic variations at particular loci (such as maturity) could be selected. An allele-focused approach could be replaced or augmented by a genomic prediction approach to predict the breeding value of each accession held in the collection (Meuwissen *et al.* 2001; Habier *et al.* 2007; VanRaden 2008). Such predictions on breeding value, especially when compared to some well-known adapted checks, would greatly increase the value of germplasm collections by giving breeders a means to identify those accessions (of the thousands that are available) meriting their attention (Longin and Reif 2014).

The USDA Soybean Collection dates back to 1895, with record keeping formally starting in 1898. A large share of the accessions (~5,000) were collected as part of the expedition of P.H.

Dorsett and W.J. Morse in Asia between 1924 and 1932 (Nelson 2011). The USDA Soybean Germplasm Collection (hereafter referred to as the Collection) is one of the most intensely used germplasm collections in the world, and the most intensely used in the NPGS (Nelson 2011). Remarkably, the entire collection has been genotyped with 50K SNPs (Song *et al.* 2015), creating a tremendous resource for understanding the distribution of genomic variation in the Collection and how it relates to phenotypic variation.

We assessed the usefulness of the genomic and phenotypic data collected on 9,171 records from the Collection for developing genomic prediction models to evaluate the genetic value of accessions held in the collection for the complex, yet economically important, traits of protein, oil, and yield. Moreover, we investigated factors affecting prediction accuracy such as training set composition both in terms of subpopulation membership and trial locations. Our results are the first report on using comprehensive, extensive data gathered over time by the curators of a germplasm collection for making genomic predictions that will help breeders select accessions in a more rational manner.

## Materials and Methods

### Phenotypic and genotypic data

The USDA Soybean Germplasm Collection contains approximately 18,500 accessions of *Glycine max.* The phenotypic data used in this study was obtained from the USDA Soybean Germplasm Collection evaluations conducted periodically to characterize newly acquired accessions for basic morphological, agronomic (including yield), and seed quality traits. Data from 25 trials were analyzed (Table 1). Dates of the data sets range from 1963 to 2003 and locations include Urbana, IL; St. Paul, MN; Lexington, KY; and Stoneville, MS. The majority (5,731) of accessions were evaluated in just one trial, 2,976 accessions were evaluated in two trials, 50 accessions were evaluated in three trials, 11 accessions were evaluated in four trials, and three accessions in five trials. Accessions originally classified as maturity group 0 were mostly evaluated in St. Paul with a small number evaluated in Urbana. Classification of these accessions has now been refined to include 00 and 000 classification. Maturity groups I – III were predominantly evaluated in Urbana with some MG I and II evaluated in St. Paul. MG IV were evaluated in Lexington, Urbana and, to a small extent, Stoneville. Accessions belonging to MGs V – IX were evaluated in Stoneville with the exception of seven MG V accessions being evaluated in Urbana in 2001–02 (Table 1). All trials were blocked by MG. The 1MN63 and 1IL64 trials included two replications planted within the same year. All other trials also included two replicates, but replicates were planted in two separate years. Field plots comprising the trials conducted between 1963 and 1966 were two rows per entry, 2.4 m long and 1 m apart, except for 1MN63 in which row spacing was 0.90 cm. Starting in 1980, trials consisted of four-row plots to minimize competition effects. Rows were 3 m long and 0.75 m apart at planting and end-trimmed to 2.4 m long. The only exceptions were the 1989–90 trials in St. Paul and Urbana, where rows were planted to be 4.7 m long but later trimmed to 3.2 m. Data were collected only on the center two rows.

**Table 1.**
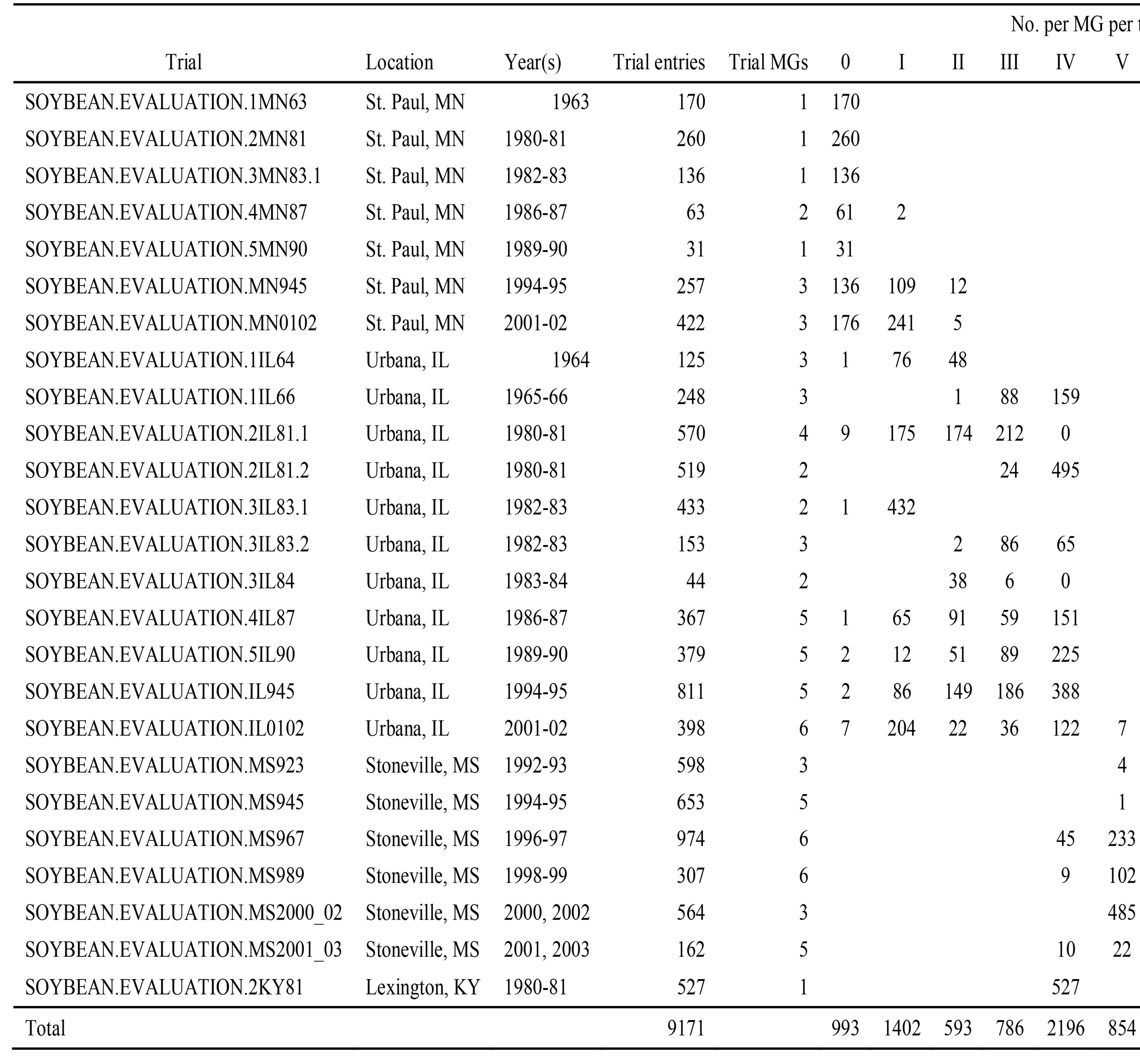
List of trials including accessions from the USDA Soybean Germplasm Collection, location of trial, year of trial, number of accession entries in each trial, number of maturity groups (MGs) in each trial, and distribution of accessions among MGs in each trial.

Protein and oil were also measured using seeds of accessions stored in cold room of the Urbana maintained Collection. This dataset is named S0YBEAN.CHEMICAL.NB.2009 and consists of 2721 samples. Soybean samples were sent from the Collection to St. Paul, MN where they were ground and scanned by NIR (Foss 6500) at the University of Minnesota. All accessions included in this set were also grown and phenotyped as part of other trials.

The traits analyzed for this study were seed yield, oil, protein, lodging, and early shattering. Seed yield was measured as the machine harvestable seed weight per plot adjusted to 13% seed moisture and expressed as Mg ha^−1^. From 1963 – 1966 protein concentration was determined using the Kjeldahl method and oil concentration was determined using the Butt extraction method. From 1981 and beyond, oil and protein concentration were determined using near-infrared reflectance on ground samples. Lodging is rated on a 1-5 scale with one given to plots with 100% erect plants and 5 given to plots with prostrate plants. Early shattering is scored at harvest on a 1–5 scale, where 1 = no shattering, 2 = 1 to 10% shattering, 3 = 10 – 25% shattering, 4 = 25 – 50% shattering, and 5 = greater than 50% shattering. More detailed trait descriptions and information on methods of measurement can be found at http://www.ars-grin.gov/cgi-bin/npgs/html/desclist.pl?51.

In addition to the phenotypic data routinely collected by the USDA and collaborators on the collection, an independent data set on MGs I-V PIs was obtained to serve as an additional validation set. These data were collected by co-author J.E. Specht at the University of Nebraska in 2003 and 2004. Briefly, 101 accessions were selected from a larger set of approximately 1500 accessions on the basis of acceptable lodging, seed shattering, disease resistance, and overall appearance. Most of these 101 accessions belong to MGs II and III. They were evaluated in field trials under two water regimes, dryland and full irrigation, at Lincoln, NE. Plots were arranged in a randomized complete block design with four replications per water regime. Replications receiving the same water treatment were blocked together in the field. Plots consisted of two rows 0.76 m apart and 2.90 m long. Plots were machine harvested and seed yield was adjusted to 13% seed moisture. Protein and oil concentration were measured using near-infrared reflectance spectroscopy. For use here, the data were divided into four water regime-year combinations. A linear model was fit to each dataset separately to calculate estimates of broad-sense heritability on an entry-mean basis. The linear model included rep (fixed) and accession (random).

The original genotype data set consisted of 52,041 single nucleotide polymorphisms (SNPs) scored using the Illumina Infinium SoySNP50K BeadChip as described by (Song *et al.* 2013). The SNP data is publicly available at http://www.soybase.org/dlpages/index.php. SNPs with greater than 80% missing scores and minor-allele frequencies less than 0.01 were removed from the data set, leaving 38,452 SNPs for analysis and genomic prediction model training.

### Subpopulation assignment

The effect of predicting across and within subpopulations was investigated. Previous research found that country of origin and MG explain only a small proportion of the subpopulation structure (Bandillo *et al.* 2015). Accessions were clustered using ADMIXTURE (Alexander *et al.* 2009) to objectively assign accessions to more genetically differentiated subpopulations. ADMIXTURE provides model-based estimations of ancestry based on multilocus genotype data. A number of subpopulations, K, is defined by the user. Each individual is assigned a membership probability to each subpopulation. For this study, the conversion from membership probabilities to discrete subpopulation memberships was accomplished by assigning each accession to the subpopulation which it had the highest probability of belonging to. Determining the value of K was accomplished using the differences from the estimated 10-fold cross-validation errors (CV) obtained from ADMIXTURE for successive K-values (ΔCV). Although the election of an optimal number of subpopulations is not a critical objective of this research, the K value at which ΔCV plateaued was chosen.

### Models

The Bayesian models here presented include genetic and non-genetic (or structural) covariates. The non-genetic covariates were included to remove, as much possible, the phenotypic variance generated by environmental and population structural factors such as location and maturity group. Since all models have the same linear predictor form, at this point, only the general structure is shown and further specifications will be given to stress differences among models.

The linear predictor can be written as 
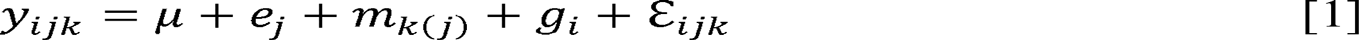
where *μ* is the overall mean common to all phenotypes, *e_j_* is the effect of the *j^th^* trial (for j=1,…,26); m_k_(j)) is the effect of the k^th^ maturity group nested in the *j^th^* trial; *g_i_* is the additive genetic effect of the i^th^ accession modeled using whole-genome markers; and *ε_ijk_* is the residual. Residuals are assumed to be independent and identically distributed (IID) following a normal distribution with mean zero and variance 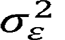. Since the effects of the maturity group are expected to change in accordance with the latitude of the trial locations, these effects were considered as nested within trials. Flat priors were given to the trial and maturity group effects to approximate fixed effects in maximum likelihood estimation.

The additive genetic effect of the i^th^ accession is modeled as a linear combination of random marker effects represented by 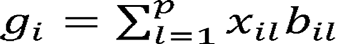, where *p* is the number of markers, *x_il_* is indicator variable for the l^th^ marker scored on the i^th^ accession with *b_il_* being the marker effect. The election of the prior distribution of the random terms enables the model to perform different actions with respect to the treatment of these marker effects as described below. A comprehensive review of the five popular models used for genomic selection (GS) can be found in (de los Campos *et al.* 2013), but a very brief description follows:.

### Genomic Best Linear Unbiased Prediction (G-BLUP)

A convenient re-parameterization to reduce the computational burden is given by considering **g** = **Xb** with **g** = *{g_i_}.* From the properties of the multivariate normal distribution 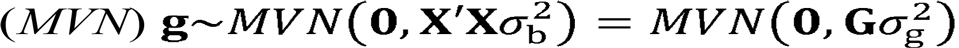 where **G** = *{G*_ii′_} an *n×n* symmetric matrix whose entries are given by 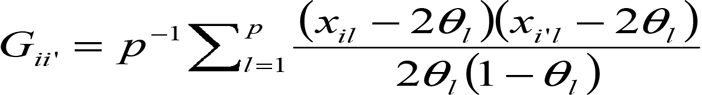 and θ*_l_* is the estimated allele frequency at the *l*^th^ marker. This matrix is known as the genomic relationship matrix (GRM) whose entries describe genomic similarities among pairs of accessions. The posterior mean of **g** is the best linear unbiased predictor of **g**, 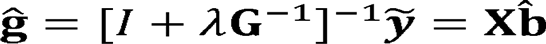, where 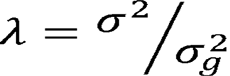is obtained via restricted maximum likelihood (REML) methods.

### Bayesian Least Absolute Angle and Selection Operator (LASSO)

The structure of this model is very similar to that from the *Bayes A* model; however, the marker-specific prior variances are assumed IID exponential, 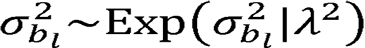where λ in this case is a regularization parameter that controls the shape of the exponential prior distribution.

The marginal prior distribution of the marker effects becomes a double exponential distribution (DE).

### Bayes B

Bayes B is a variable selection model allowing some proportion (7T) of marker effects to be null and the remaining (1-π) to be non-null. This is captured with a mixture density: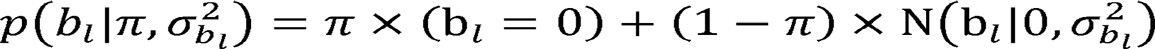 πwith as the proportion of markers with null effect. With this consideration, the marker-specific prior distributions of the non-null marker effects are a scaled inverted chi-squared distribution,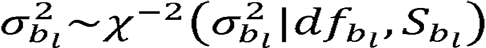. The prior distribution of the marker-specific variance parameter for the non-null proportion of marker effects is similar to the one used in Bayes A. To completely specify this model, a beta prior distribution is assumed for the proportion parameter such that π~Beta(*P*_0_,π_0_)with *p*_0_ > 0 and π_0_ϵ[0,l]. All of these assumptions result in the marker effects having a marginal prior distribution comprised of an IID mixture of a point mass at zero and a scaled-*t* distribution.

Hyperparameters for all models were set using the rules described in (de los Campos *et al.* 2013). All models were implemented using the BGLR package (Pérez and de los Campos 2014).

### Cross-Validation Schemes

A series of cross-validation (CV) schemes was designed to assess the usefulness of genomic predictions for selecting accessions as well as optimizing the construction of genomic prediction training sets. To accomplish the latter goal, several different grouping criteria for splitting the data were used in order to create variable training-testing relationships. The first grouping criteria involved splitting the data by trial (i.e., 26 trials for oil and protein; 25 trials for yield). A second grouping criteria used genetic criteria to split the entire population of accessions into nine subpopulations as described above. Finally, the training-testing sets were grouped by geographical location, which in this case was defined by state in which the evaluation trials were conducted (i.e., MN, IL, or MS). The KY data was dropped from this analysis since only one trial was conducted in KY.

Four CV schemes were applied to each grouping criteria. Each CV scheme mimicked the problem of predicting accessions without data. To accomplish this, all phenotypic records of any accession in the validation set was removed from the training set before model training. Each CV scheme is described individually.

*Leave-one-accession-out within groups (One/Group):* Within each group (i.e., trial, subpopulation, state) each accession is predicted, one at a time, using as the training set the data from the remaining accessions in the same group. This procedure is repeated until all accessions in the group are predicted. To assess predictive ability, observations and predictions are compiled and correlated for each group separately.

*Leave-one-accession-out across groups (One/All):* This CV is the same as *One/Group* except training sets consists of data from all groups rather than only a single group.

*Leave-one-group-out (Group/All):* Here, each group is predicted using a training set consisting of data from the other groups only. The training set does not include data from the group comprising the validation set.

*Group-by-group (Group/Group):* A whole group is predicted using the information from another, single group. This procedure is repeated for all possible combinations.

A schematic of these cross validation schemes is displayed in Figure 1.

Predictive ability was assessed using Pearson’s product-moment correlation coefficient on the vectors of genomic predictions and observed phenotypes adjusted for trial and MG effects. Confidence intervals were computed using the bootstrap procedure with 10,000 bootstrap replicates.

**Figure 1.**
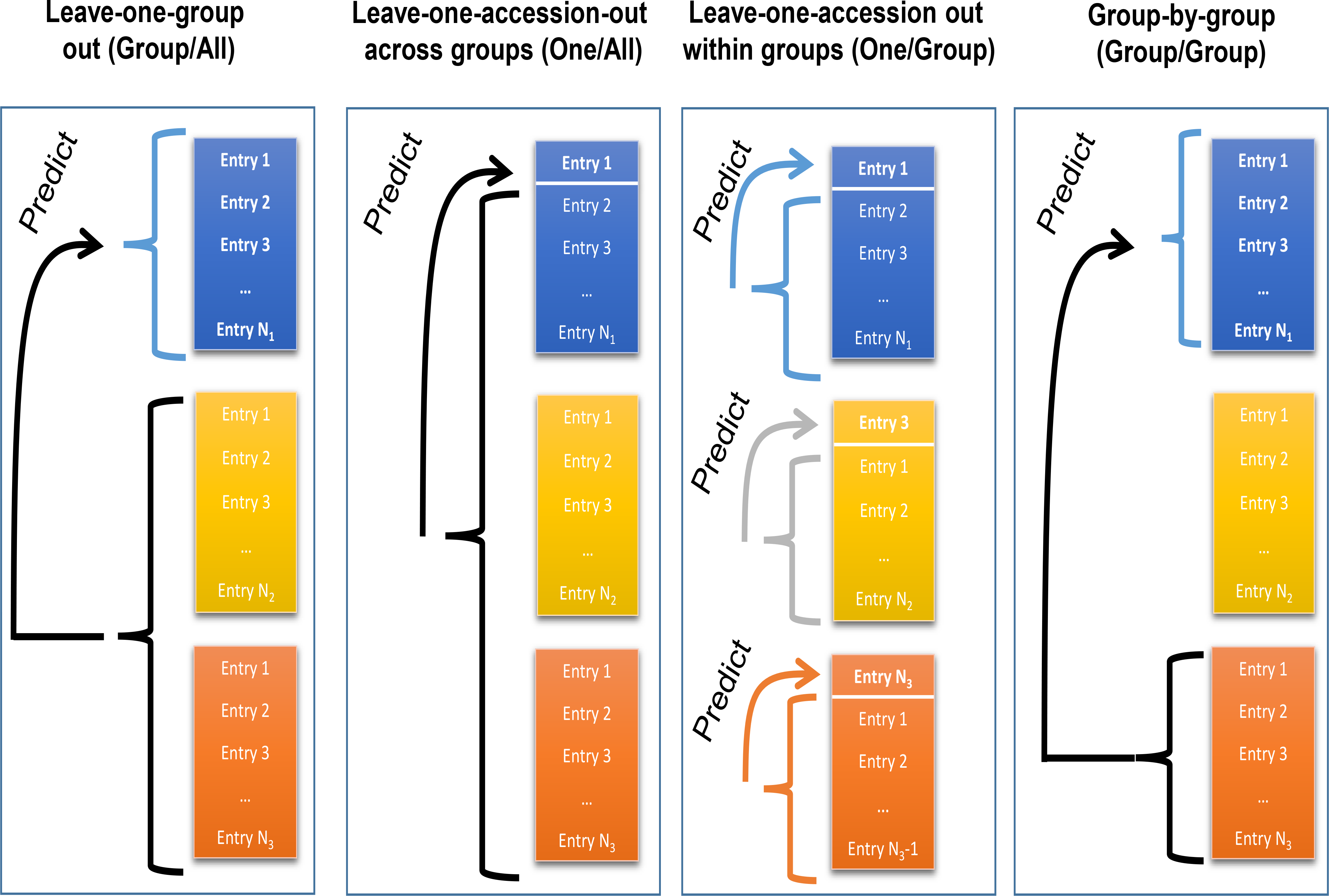
A diagram of the four cross validation schemes used to validation genomic predictions. Each of the colors represents a different group. Groups are comprised of trials, states, or genetic clusters (see Methods section). Arrows point from the training set to the validation set.

## Results

An initial assessment of predictive ability for oil, protein, and yield was made by performing *Group/All* CV and evaluating predictive abilities among the 25-26 trials with the five models described above. Average predictive abilities were moderate to very high for most trial and trait combinations (Figure 2; Supplementary Table 1). For oil, predictive ability ranged from 0.46 to 0.92 with a median across trials of 0.69. Predictive abilities for protein were lower, especially on the low end, ranging from 0.29 to 0.92 with a median of 0.56. Genomic prediction for accession yield, typically the most difficult trait to predict, was more successful than expected, yet highly variable, ranging from 0.17 to 0.79. The median predictive ability for yield was 0.64. Predictive abilities among the five models were compared for each trial and trait combination. Very little to no difference among models was observably evident. This can be seen by examining the performance of three representative models (G-BLUP, Bayes B, Bayes LASSO) in Figure 2. For this reason, G-BLUP was exclusively used for all subsequent analyses.

**Figure 2.**
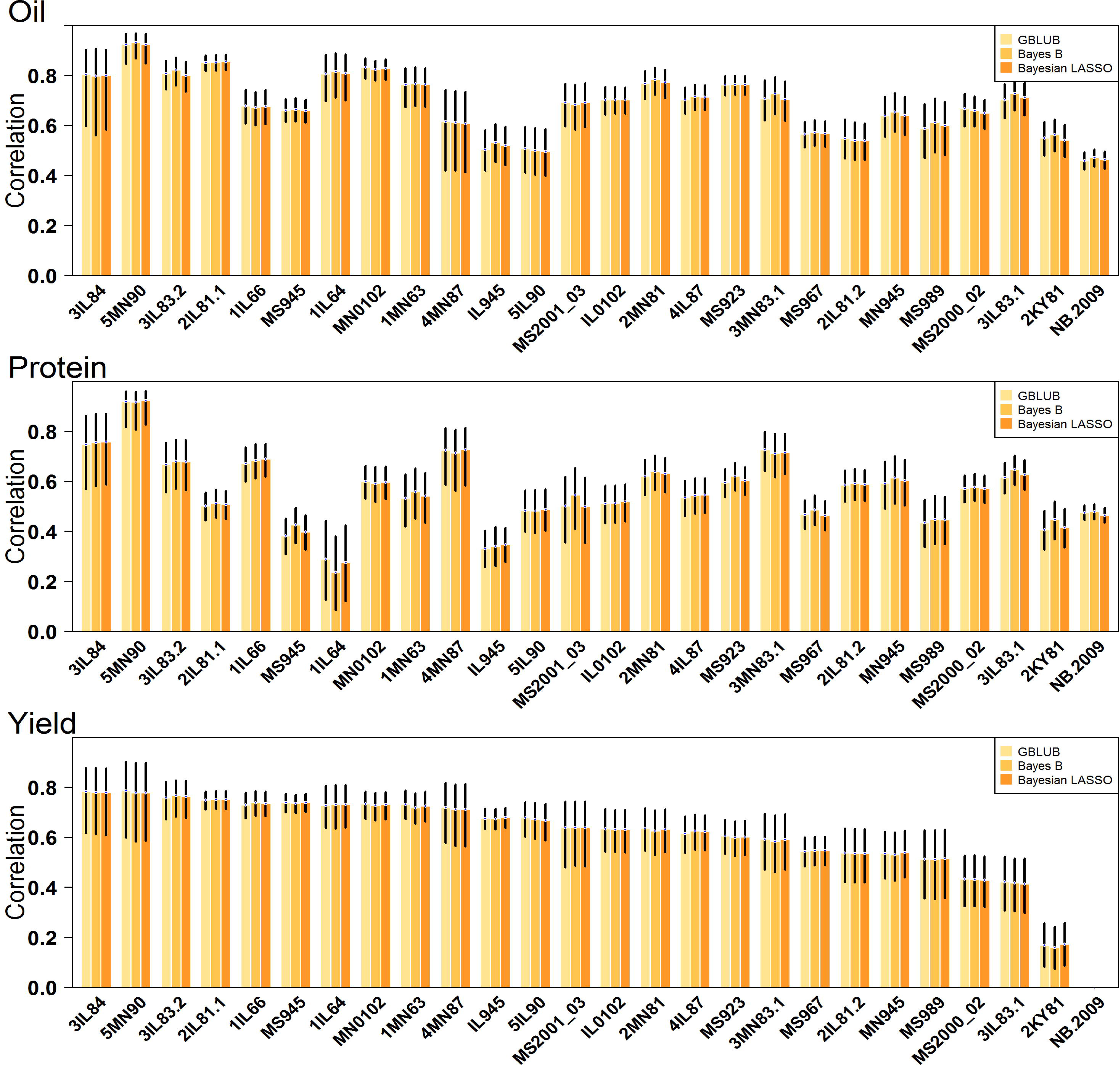
Predictive abilities for oil, protein, and seed yield for each of the 25 trials. Predictions were made using the *Group/All* cross validation scheme and three different models: genomic best linear unbiased prediction (G-BLUP), Bayes B, and Bayesian LASSO. The black bars display the 95 percent confidence intervals.

It is important to remember that the validation phenotypes were adjusted for MG effects and thus variation explained by the genomic prediction model is independent of any variation between maturity groups and the predictive abilities calculated reflect the ability to predict within maturity groups. Moreover, reported predictive abilities are correlations between predictions and phenotypes corrected for MG, with no adjustment being made for the heritability of the validation phenotypes. Since the validation phenotypes are imperfect estimates of true additive genetic values, the reported predictive abilities are likely downwardly biased estimates of the true prediction accuracies being defined as the correlation between the predictions and true breeding values.

Since most of the accessions analyzed were unimproved landraces, an important consideration is the degree to which variation in lodging and shattering influence variation in machine harvestable seed yield. Shattering is a genetically simpler trait compared to yield (Funatsuki et al., 2014), and genomic prediction models trained using data from landraces might be simply predicting shattering rather than purely seed yield. An analysis of the phenotypic data did reveal that machine harvestable grain yield was negatively correlated with shattering and lodging, with mean correlation coefficients being −0.27 and −0.21, respectively (data not shown). In order to eliminate the influence of shattering and lodging on variation in seed yield, shattering and lodging scores were fit as fixed covariates both in the G-BLUP model and to calculate adjusted seed yield phenotypes for validation. Predictive ability was calculated as it was for Figure 2 using the 8517 records with available shattering and lodging scores. Predictive abilities for yield were reduced as expected when variation for lodging, shattering, or both was removed through the use of covariates (Supplementary Table 2). The average reduction in seed yield predictive ability, expressed as a percentage of the original predictive ability, was near 10% when either lodging or shattering were accounted for (Supplementary Table 2). When both traits were fit as covariates, the reduction in predictive ability was 23% on average across trials (Supplementary Table 2). Predictive ability was not reduced at all, or very little, in some trials, whereas in others it was reduced by as much as 47%, indicating shattering and lodging affected seed yield to different degrees across trials. A similar analysis was performed on maturity date, but maturity date had a negligible effect on seed yield predictions after correction for MG effects (results not shown).

The value of training genomic prediction models for prediction of accession performance was further evaluated by using an independent set of MG II and III accessions evaluated in multiple environments, each with four replications, for the measured traits of oil, protein, and seed yield. Entry-mean heritabilities were high due to the highly replicated design, ranging from 0.83 on average for yield to 0.91 on average for protein and oil. Genomic predictive abilities, on average, were 0.58–0.59 for protein and oil and 0.49 for yield (Table 2). These values are somewhat lower than the predictive abilities estimated for the GRIN trials, perhaps because the Nebraska trials included less genetic variability because accessions were pre-selected on the basis of agronomic performance. They were, however, similar to the predictive abilities for seed yield in the GRIN trials when seed yield values were adjusted for shattering and lodging. This result indicates that the genomic prediction models can still discriminate among relatively poor and good performing accessions within sets previously selected for agronomic performance. A comparison was made between the genomic predictive ability and correlations between the data available from the GRIN trials and the phenotypes collected in the highly replicated Lincoln, NE trials. Genomic predictive ability was consistently better than the GRIN phenotypes for protein and oil, although the confidence intervals did overlap (Table 2). For yield, the two methods were similar for three of the four Lincoln, NE trials, and genomic prediction was numerically better than the phenotypic data in the fourth comparison (DRY-2003; Table 2). This result suggests that the genomic predictions are at least as good as the phenotypic data in GRIN, and thus it may be a useful tool for choosing among the non-phenotyped accessions held in the collection or newly collected accessions.

**Table 2.**
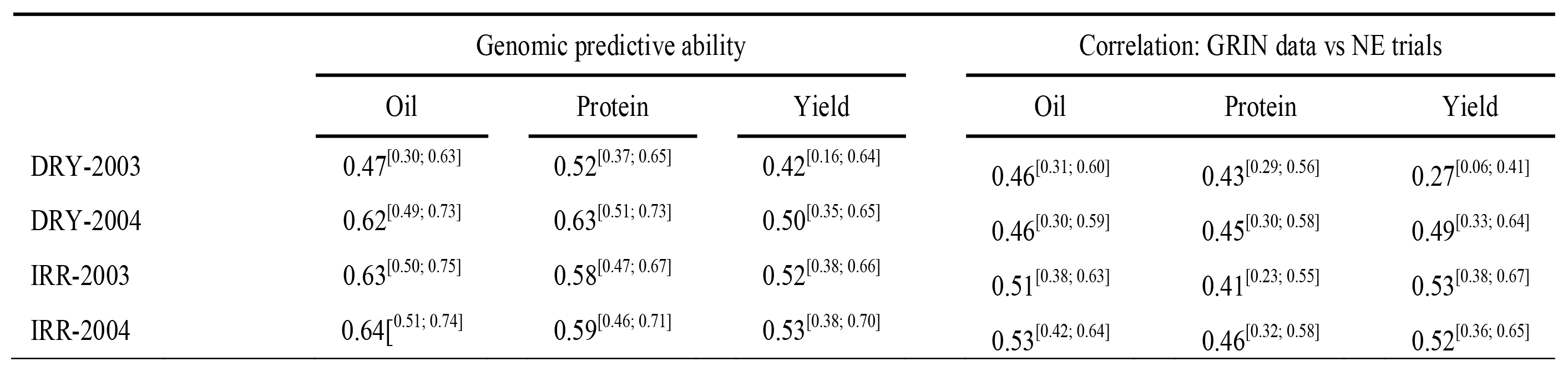
Genomic predictive abilities using the Lincoln, NE data as validation data, and correlations between the phenotypic data available in GRIN and the Lincoln, NE data. Each year two trials were conducted with two water regimes, dryland (DRY) and irrigated (IRR). Results for oil, protein, and yield are displayed.

|  | Genomic predictive ability |  |  | Correlation: GRIN data vs NE trials |  |  |
| --- | --- | --- | --- | --- | --- | --- |
|  | Oil | Protein | Yield | Oil | Protein | Yield |
| DRY-2003 | 0.47<sup>[0.30; 0.63]</sup> | 0.52<sup>[0.37; 0.65]</sup> | 0.42<sup>[0.16; 0.64]</sup> | 0.46<sup>[0.31; 0.60]</sup> | 0.43<sup>[0.29; 0.56]</sup> | 0.27<sup>[0.06; 0.41]</sup> |
| DRY-2004 | 0.62<sup>[0.49; 0.73]</sup> | 0.63<sup>[0.51; 0.73]</sup> | 0.50<sup>[0.35; 0.65]</sup> | 0.46<sup>[0.30; 0.59]</sup> | 0.45<sup>[0.30; 0.58]</sup> | 0.49<sup>[0.33; 0.64]</sup> |
| IRR-2003 | 0.63<sup>[0.50; 0.75]</sup> | 0.58<sup>[0.47; 0.67]</sup> | 0.52<sup>[0.38; 0.66]</sup> | 0.51<sup>[0.38; 0.63]</sup> | 0.41<sup>[0.23; 0.55]</sup> | 0.53<sup>[0.38; 0.67]</sup> |
| IRR-2004 | 0.64<sup>[0.51; 0.74]</sup> | 0.59<sup>[0.46; 0.71]</sup> | 0.53<sup>[0.38; 0.70]</sup> | 0.53<sup>[0.42; 0.64]</sup> | 0.46<sup>[0.32; 0.58]</sup> | 0.52<sup>[0.36; 0.65]</sup> |

A key question when using predictions for accession selection relates to the enrichment of the selected set. While correlations are a good indicator of how successful genomic predictions could be used for this purpose, we desired to directly look at this by calculating the frequency of “selected” accessions observed to be better than the mean or in the bottom 10% based on actual field trial data. The top 10% of accessions were chosen on the basis of their genomic predictions using G-BLUP and *Group/All* as described above. Shattering and lodging were not adjusted for in this analysis as we assumed breeders would want select accessions with high machine harvestable seed yield. We found that, on average, a high percentage of accessions among the top 10% based on predictions were observed to be better than the trial mean. This value was 89% for oil, 80% for protein, and 88% for yield (Table 3). In the case of yield, 100% of the selected accessions were observed to be better than the population mean in five trials. Another key outcome would be the avoidance of poorly performing accessions. The top 10% based on predictions very rarely included accessions observed to be in the bottom 10%. The average observed frequency across trials was only 0 – 2% depending on the trait (Table 3). A frequency of 0% was observed for more than half the trials. This result indicates that predictions can very effectively eliminate the worst performing accessions.

**Table 3.**
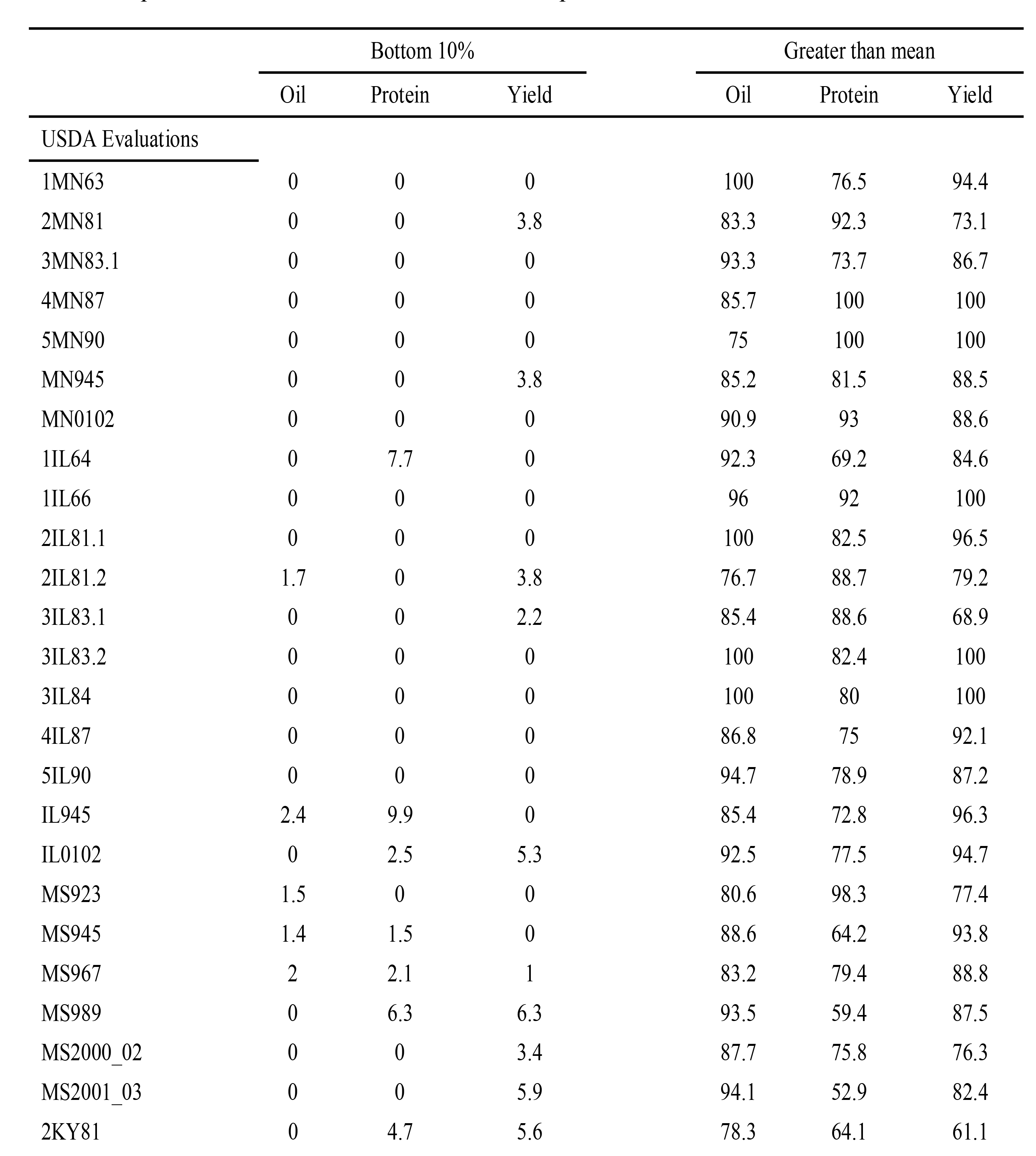
Tabled values are percentages of accessions among the top 10% of accessions based on predictions that were observed to be in the bottom 10% or greater than the mean based on phenotypic data from each listed trial. Data for both the USDA Soybean Germplasm Collection evaluations and J.E. Specht trials conducted in Lincoln, NE are presented.

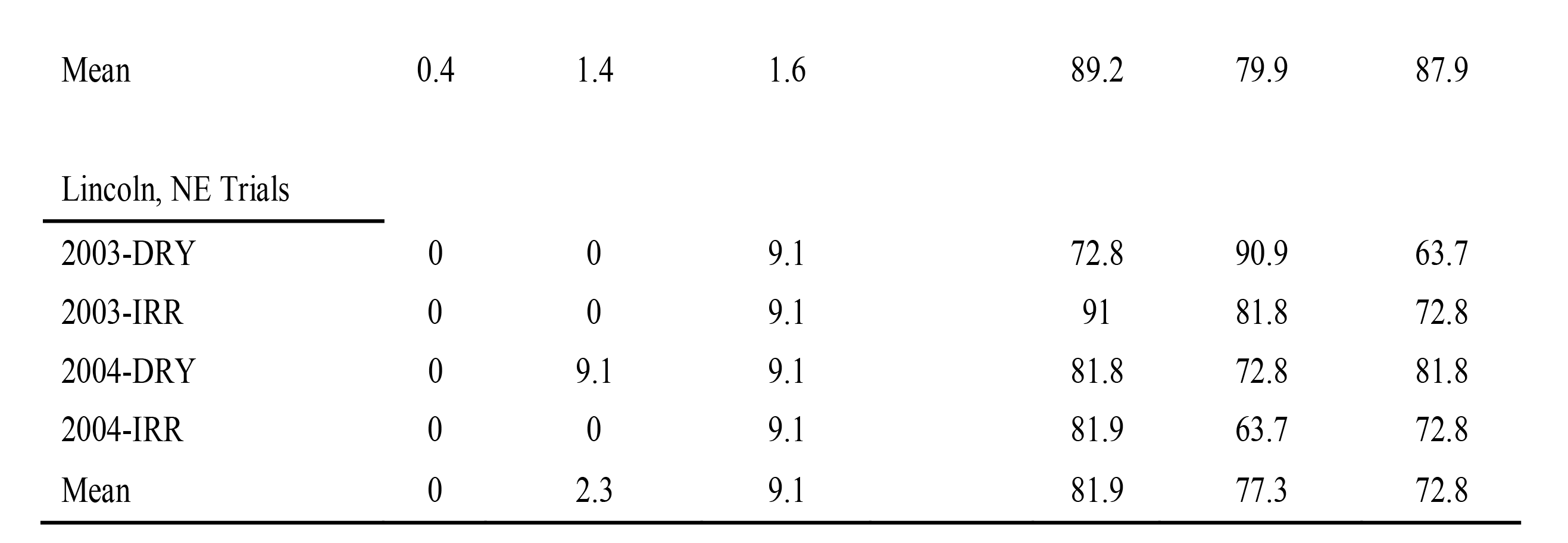

## Trials

The effect of combining *versus* separating trial data was evaluated by performing within-trial cross validation (One/Group), between-trial cross validation *(Group/All),* and combining data across all trials *(One/All)* (Figure 1).

Within-trial predictive abilities were moderate to high for all traits, being greater than 0.60 in most cases for oil and yield (Figure 3; Supplementary Table 1). Predictive abilities were slightly lower for protein. Only a very subtle improvement was observed when data across all other trials was added to the training set *(One/All),* with differences ranging from 0.04 (protein) to −0.01 (yield) (Supplementary Table 1). While on average there was very little difference, the range across trials was considerable, and it appeared that there were benefits to including all data in the extreme cases. In the case of the 3IL84 trial, for example, it was observed that predictive ability could be increased from 0.31 to 0.81 for oil and 0.44 to 0.75 for protein when data was combined across all trials compared to a within-trial training set only. On the negative side, we observed that predictive ability was decreased by 0.05 for oil and 0.06 for protein, in the case of IL66 and IL945 trials, respectively. It appeared that for protein and oil, benefits to combining across trials were much more dramatic compared to any reductions in predictive ability (Figure 3; Supplementary Table 1). The differences between *One/Group* and *One/All* were more uniformly distributed in the case of yield, with a reduction of 0.06 for MN945 and a gain of 0.14 for 2KY81.

**Figure 3.**
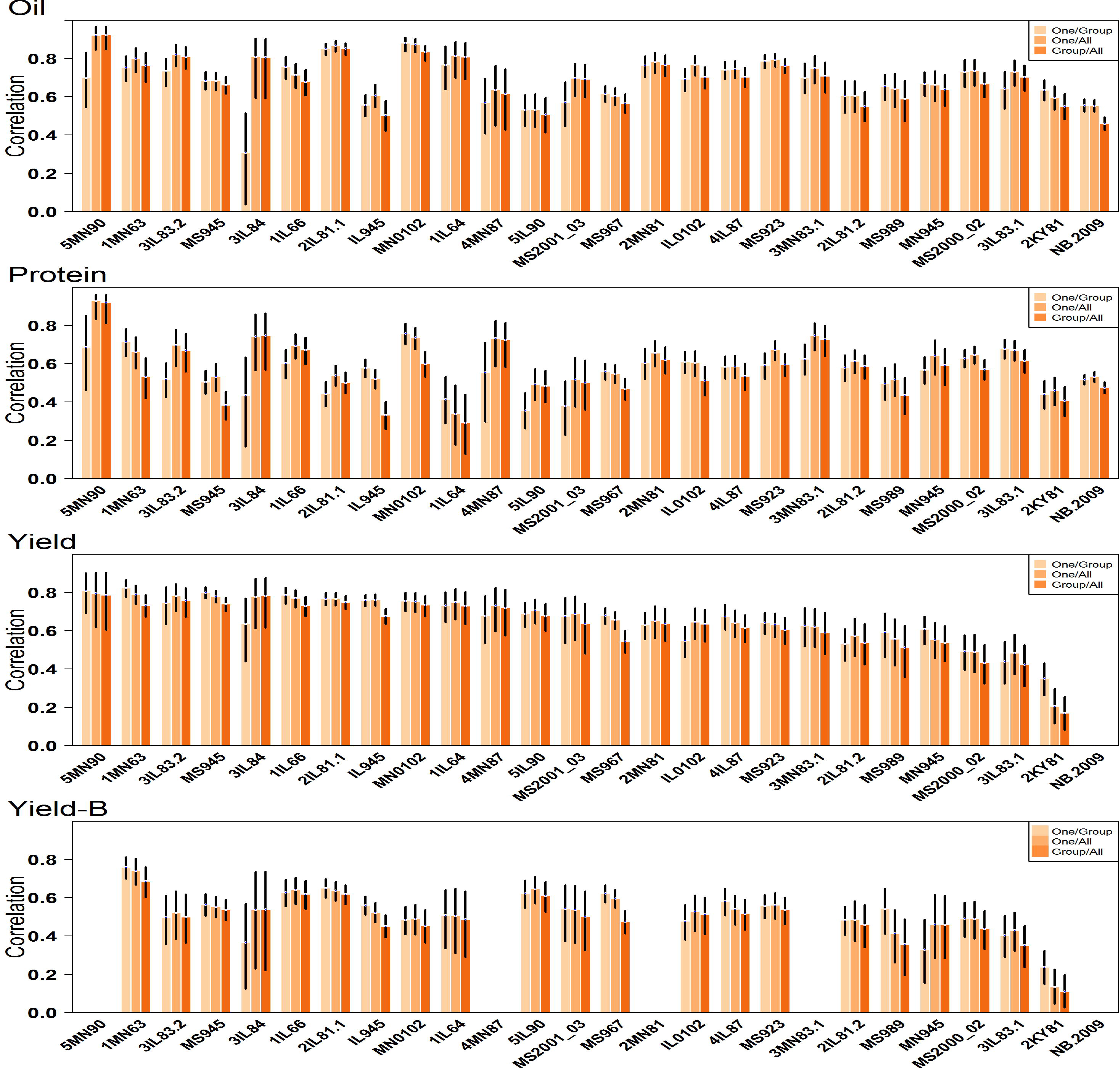
Predictive abilities for oil, protein, and seed yield for each of the 25 trials. Predictions were made using the *One/Group, One/All,* and *Group/All* cross validation schemes. The black bars display the 95 percent confidence intervals.

Using data from the same trial(s) in both training and validation sets creates the unrealistic advantage of including the trial-specific GxE effects contained in the validation data. Because the exact same environmental conditions specific to individual trials would not be observed again, a better assessment of the usefulness of these GRIN training sets for predicting future trial performance would be attained using the *Group/All* CV. The *Group/All* CV correlations were very close, on average, to the *One/Group* and *One/All* CV correlations (Figure 3; Supplementary Table 1), indicating that the sheer volume of data can overcome any lack of shared GxE effects.

A trial-by-trial CV *(Group/Group)* results in highly variable predictive abilities. In many cases, the predictive abilities between trials was zero, but in some cases, the predictive ability reached as high as 0.90 (e.g., oil, 2MN81 predict 5MN90) (Supplementary Table 3). The average predictive ability for the *Group/Group* CV was 0.49 for oil, 0.30 for protein, and 0.45 for yield, which is far less than the predictive abilities observed using *Group/All* CV. This illustrates the expected advantage of combining data across many trials to form a training set.

By ordering the trials by state, it is apparent that the northern locations of MN and IL predicted one another relatively well as compared to be predictive ability between MS and the northern locations. This pattern was more prominent for yield (Supplementary Table 3).

## States

Given the pattern observed when predicting between trials conducted in different states, we desired to look at this more closely by setting up a CV based on trial geographical location. The distribution of data points across states is as follows: 4,047 records from 11 IL trials; 1,339 records from 7 MN trials; 3,258 records from 6 MS trials (Table 1). The MN trials consisted of only MG 0 - II accessions with the majority (72%) belonging to MG 0. The IL trials were predominantly comprised of accessions from MGs I - IV, with less than 1% being from MGs 0 and V. The MS trials only consisted of accessions from MGs IV - IX.

As expected, a training set including data from the state being predicted *(One/Group* or *One/All)* performed much better than training sets not including data from the state being predicted *(Group/All)* (Figure 4; Table 4). A key question we wanted to address with this analysis was whether training sets should be created by dividing data among states, or if a universal training set including all data - regardless of state - would perform just as well or better. Little to know differences were observed between these two CV schemes for any trait and state combination (Figure 4; Table 4). This finding suggests that predictive abilities are not improved by maximizing training set size by combining across states, nor are they reduced by including data from environments as different as MS when predicting relative performance of early MG accessions in MN. A similar pattern was observed when variation for lodging and shattering was removed through inclusion of covariates.

**Table 4.** Predictive abilities for oil, protein, and yield estimated using State as the grouping factor. *One/Group* estimates are on the diagonal and *Group/Group* estimates are on the off-diagonal.

|  | <u>MN</u> | <u>IL</u> | <u>MS</u> |
| --- | --- | --- | --- |
| MN | 0.80 | 0.71 | 0.53 |
| IL | 0.56 | 0.68 | 0.52 |
| MS | 0.43 | 0.51 | 0.62 |

Protein
|  | <u>MN</u> | <u>IL</u> | <u>MS</u> |
| --- | --- | --- | --- |
| MN | 0.70 | 0.43 | 0.42 |
| IL | 0.31 | 0.55 | 0.39 |
| MS | 0.21 | 0.37 | 0.52 |

Yield
|  | <u>MN</u> | <u>IL</u> | <u>MS</u> |
| --- | --- | --- | --- |
| MN | 0.69 | 0.59 | 0.44 |
| IL | 0.51 | 0.68 | 0.43 |
| MS | 0.40 | 0.45 | 0.66 |

**Figure 4.**
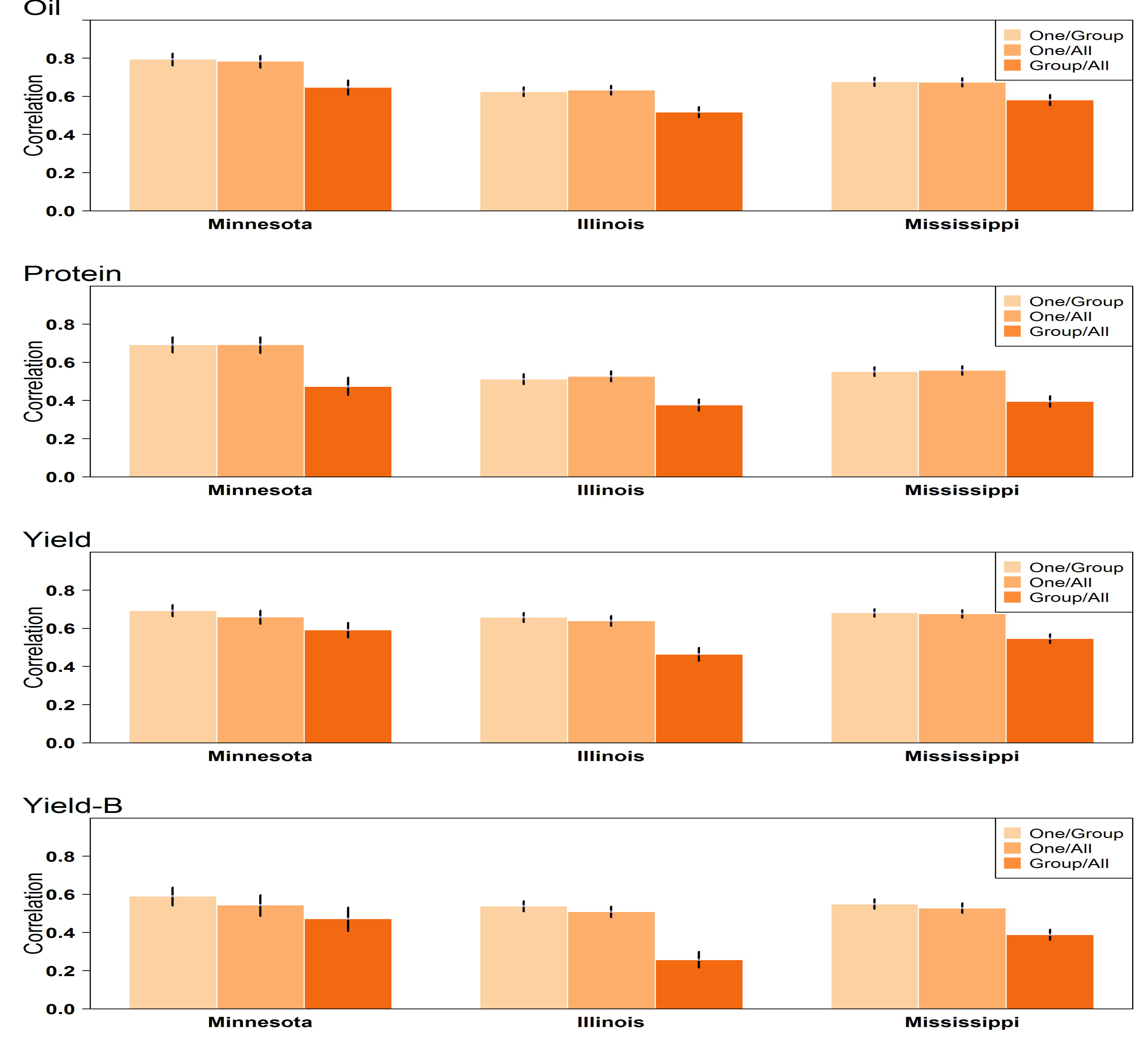
Predictive abilities for oil, protein, and seed yield for each of the three states. Predictions were made using the *One/Group, One/All,* and *Group/All* cross validation schemes. The black bars display the 95 percent confidence intervals.

## Clusters

The ADMIXTURE analysis suggested the presence of nine genetic clusters within the population of accessions used for this study (Figure S1). A visual inspection of the principal component analysis plot displayed in Figure 5 suggests that the diversity *within* clusters varies and structure *among* the clusters exists, with some clusters being more closely related than other clusters. The proximity of clusters to one another can be partially explained by MG. Clusters four, five and eight are comprised mainly of early maturity groups (0-II), whereas early and medium MGs appear in Cluster 1 (Supplementary Table 4). Clusters 2, 3, 6, 7 and 9 belong to medium and late MGs. Most clusters include good representation of at least three MGs except for cluster 3, which is dominated by MG VIII.

**Figure 5.**
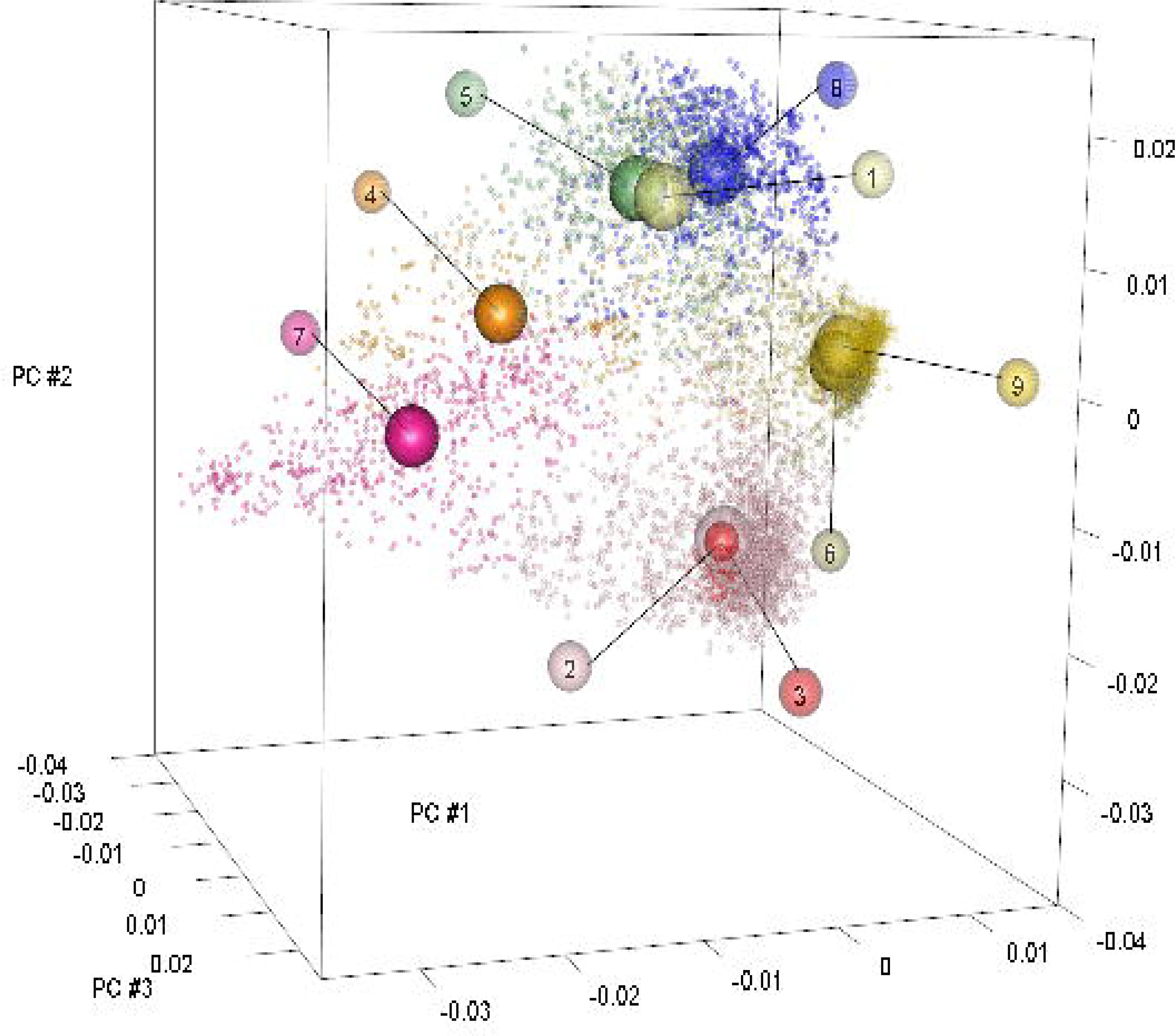
A three dimensional plot of accession values for principal components one, two, and three. The centroid of each cluster is indicated by an empty sphere. The spheres containing numbers label each centroid by its corresponding cluster designation.

In general, predictive abilities were lower for the *Group/All* scheme based on cluster compared to the *One/Group* and *One/All* schemes (Figure 6; Table 5). Without correction for shattering and lodging, the *One/All* scheme tended to produce the highest predictive abilities, although the difference between *One/Group* and *One/All* were very small. Correcting seed yield for shattering and lodging produced a different outcome where the *One/Group* scheme was markedly better for four of the nine clusters (Figure 6; Table 5). A pattern between predictive ability and relationship between clusters was not readily apparent. The only consistent result was the poor predictive ability of cluster 3, which was expected based on its limited size and variation. These results combined indicate that within-cluster information is the most valuable information. We tested whether compiling a training set by only including related clusters improved predictive abilities. To do this, clusters 1, 4, 5, and 8 were grouped, and clusters 2, 3, 6, 7, and 9 were grouped. Grouping clusters by genetic similarity did not improve predictions compared to the universal *One/All* scheme (data not shown).

**Table 5.** Predictive ability from LOAWG, LOGO, and LOAAG cross validation schemes for oil, protein, and yield using trial data grouped by genetic cluster.

| Cluster | Oil |  |  |  |  |  | Protein |  |  |  |  |  | Yield |  |  |  |  |  |
| --- | --- | --- | --- | --- | --- | --- | --- | --- | --- | --- | --- | --- | --- | --- | --- | --- | --- | --- |
|  | LOAWG |  | LOGO |  | LOAAG |  | LOAWG |  | LOGO |  | LOAAG |  | LOAWG |  | LOGO |  | LOAAG |  |
|  | Est. | 95 % CIa | Est. | 95 % CIa | Est. | 95 % CIa | Est. | 95 % CIa | Est. | 95 % CIa | Est. | 95 % CIa | Est. | 95 % CIa | Est. | 95 % CIa | Est. | 95 % CIa |
| 1 | 0.34 | [0.25; 0.42] | 0.48 | [0.40; 0.56] | 0.52 | [0.44; 0.58] | 0.40 | [0.32; 0.48] | 0.51 | [0.44; 0.57] | 0.55 | [0.49; 0.61] | 0.35 | [0.25; 0.46] | 0.43 | [0.30; 0.54] | 0.47 | [0.36; 0.57] |
| 2 | 0.54 | [0.51; 0.56] | 0.44 | [0.41; 0.47] | 0.54 | [0.51; 0.57] | 0.50 | [0.47; 0.53] | 0.36 | [0.32; 0.40] | 0.50 | [0.47; 0.53] | 0.57 | [0.53; 0.60] | 0.50 | [0.45; 0.55] | 0.58 | [0.54; 0.62] |
| 3 | 0.13 | [0.00; 0.30] | 0.32 | [0.11; 0.51] | 0.35 | [0.12; 0.54] | 0.32 | [0.17; 0.46] | 0.28 | [0.10; 0.45] | 0.27 | [0.09; 0.44] | 0.18 | [0.01; 0.35] | 0.27 | [0.11; 0.43] | 0.25 | [0.08; 0.41] |
| 4 | 0.41 | [0.30; 0.51] | 0.39 | [0.28; 0.50] | 0.45 | [0.35; 0.55] | 0.34 | [0.24; 0.43] | 0.32 | [0.21; 0.41] | 0.45 | [0.35; 0.53] | 0.44 | [0.33; 0.54] | 0.35 | [0.25; 0.45] | 0.41 | [0.31; 0.51] |
| 5 | 0.42 | [0.37; 0.47] | 0.42 | [0.37; 0.46] | 0.48 | [0.43; 0.52] | 0.46 | [0.40; 0.52] | 0.40 | [0.35; 0.44] | 0.51 | [0.46; 0.55] | 0.32 | [0.26; 0.39] | 0.38 | [0.31; 0.45] | 0.44 | [0.37; 0.50] |
| 6 | 0.52 | [0.49; 0.55] | 0.38 | [0.34; 0.42] | 0.53 | [0.49; 0.55] | 0.48 | [0.45; 0.51] | 0.30 | [0.27; 0.34] | 0.50 | [0.46; 0.52] | 0.44 | [0.40; 0.48] | 0.35 | [0.31; 0.39] | 0.45 | [0.42; 0.49] |
| 7 | 0.45 | [0.38; 0.51] | 0.4 | [0.34; 0.45] | 0.46 | [0.39; 0.52] | 0.40 | [0.34; 0.46] | 0.22 | [0.14; 0.29] | 0.37 | [0.30; 0.44] | 0.43 | [0.37; 0.49] | 0.40 | [0.34; 0.45] | 0.44 | [0.38; 0.49] |
| 8 | 0.55 | [0.50; 0.59] | 0.52 | [0.48; 0.57] | 0.58 | [0.53; 0.62] | 0.48 | [0.44; 0.52] | 0.41 | [0.38; 0.45] | 0.52 | [0.48; 0.55] | 0.45 | [0.39; 0.51] | 0.44 | [0.38; 0.49] | 0.49 | [0.43; 0.54] |
| 9 | 0.52 | [0.48; 0.56] | 0.38 | [0.32; 0.43] | 0.53 | [0.49; 0.56] | 0.44 | [0.40; 0.48] | 0.24 | [0.19; 0.30] | 0.46 | [0.42; 0.50] | 0.34 | [0.29; 0.39] | 0.22 | [0.15; 0.28] | 0.34 | [0.28; 0.39] |
| Mean | 0.43 |  | 0.42 |  | 0.50 |  | 0.42 |  | 0.34 |  | 0.46 |  | 0.39 |  | 0.37 |  | 0.43 |  |

<sup>a</sup> Obtained by Bootstrapping 10,000 the adjusted phenotypes and predicted values.

**Figure 6.**
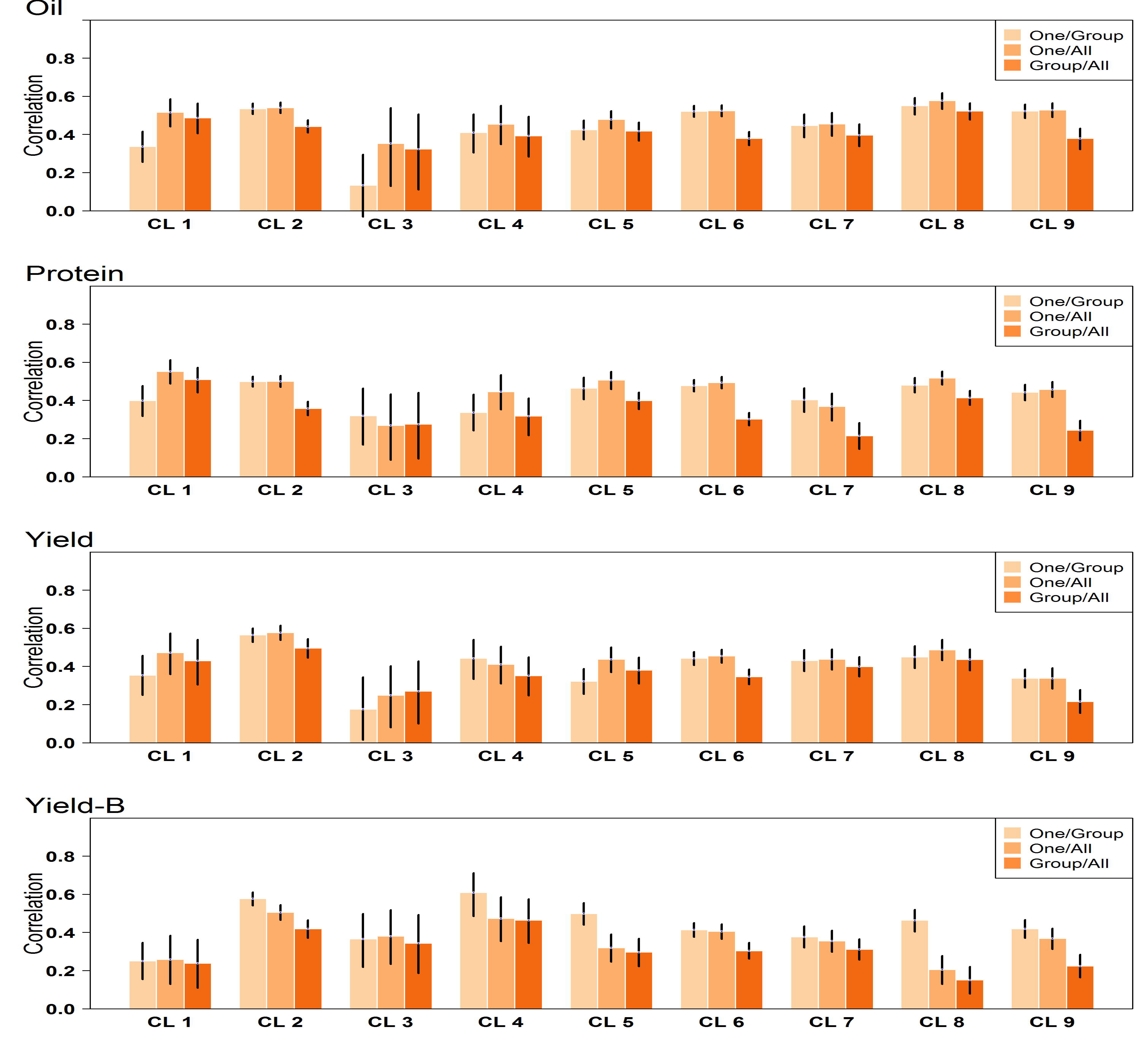
Predictive abilities for oil, protein, and seed yield for each of the nine genetic clusters (CL). Predictions were made using the *One/Group, One/All,* and *Group/All* cross validation schemes. The black bars display the 95 percent confidence intervals.

## Prediction of non-phenotyped accessions

A total of 8,771 accessions housed in the Collection have been phenotyped at least once in the 26 trials (Supplementary Table 4), but no phenotypic data was available for 7,608 accessions from GRIN when this study was designed. Genomic predictions were calculated for the non-phenotyped accessions using the full training set (i.e., all clusters, all environments) in order to assess differences in distributions between phenotyped and non-phenotyped accessions. Phenotyped accessions were predicted with the *One/All* cross-validation scheme. More specifically, we wanted to know if any non-phenotyped accessions would be predicted to be superior to the phenotyped accessions. Substantial differences were not observed with the predictions of the non-phenotyped accessions falling within the range of the phenotyped accessions (Figure 7). Nevertheless, using information in the form of genomic predictions will help breeders choose amongst those accessions that have no accompanying information, opting for those that would be expected to be above average for yield, protein, and oil and thus avoiding those accessions predicted to be inferior for these traits. Supplementary Tables 5 and 6 contain genomic predictions for the phenotyped and non-phenotyped accessions.

**Figure 7.**
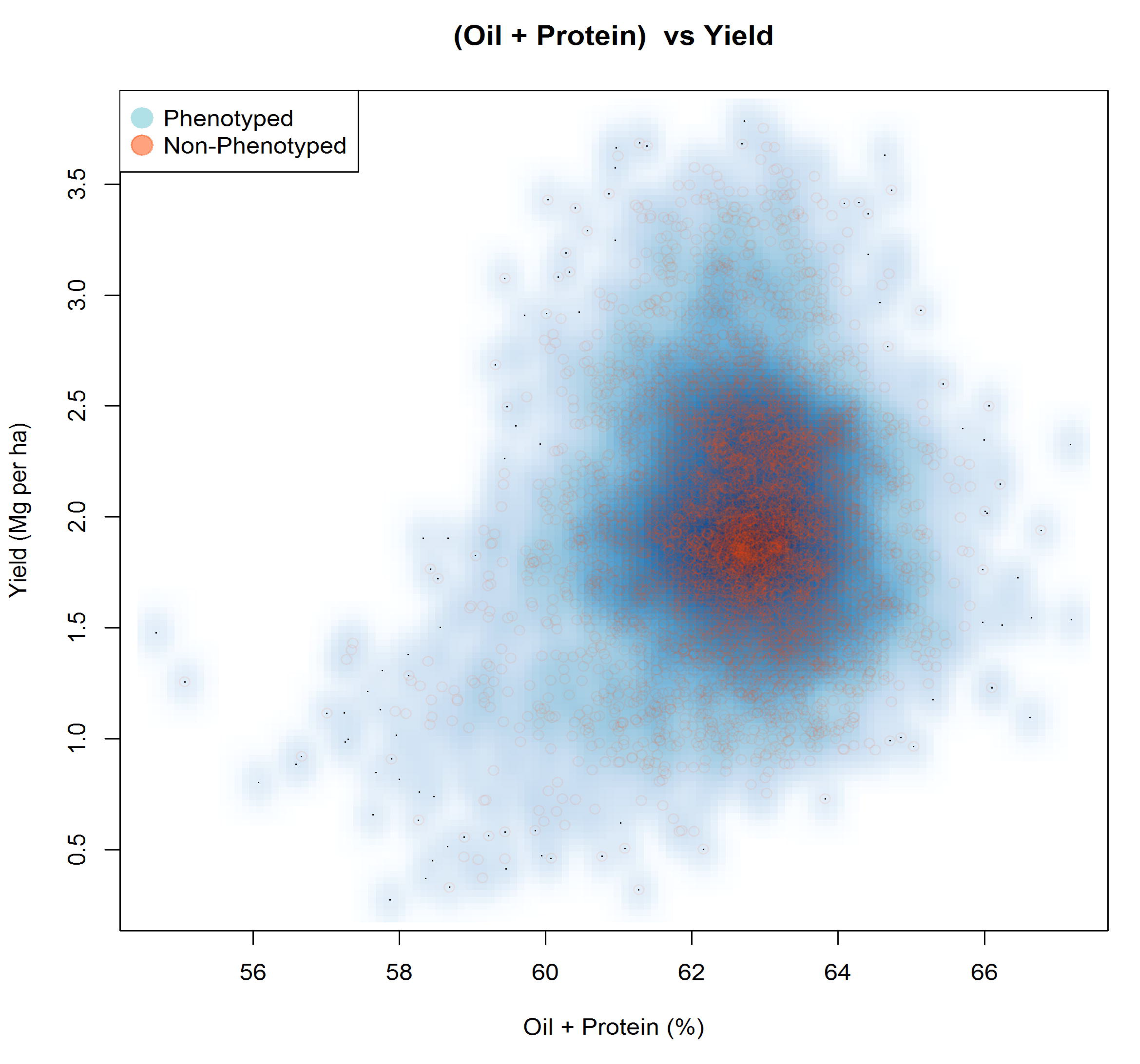
Scatter plot of genomic predictions for grain yield versus the sum of oil and protein. The intercept of each trait included in the prediction to place values back on the original trait measurement scale. Phenotyped accessions are represented by the blue density cloud, and non-phenotyped accessions are represented by the red circles.

## DISCUSSION

Crop germplasm collections hold valuable genetic diversity to help protect society against the genetic erosion of agriculturally important species for which only a limited number of genotypes are actually cultivated at any given time. It is imperative that these collections exist as dynamic, utilized sources of variation rather than as “gene morgues” as they are sometimes referred to (Hoisington *et al.* 1999). One obstacle to utilization is reliable phenotypic characterization of collections as phenotyping collections consisting of tens of thousands of accessions can be difficult and expensive. High density genotyping of entire germplasm collections, however, has become more feasible than thorough phenotyping even with the advent of phenomics platforms. This study demonstrated that historical data on accessions held in collections, when combined with high density SNP data, can be used to develop predictive models for important and complex traits of soybean. Genomic prediction models explained an appreciable amount of the variation in accession performance in independent trials, with correlations between predictions and observations reaching up to 0.92 for oil and protein and 0.79 for machine-harvestable seed yield. Predictive abilities for seed yield were reduced when variation for lodging and shattering was accounted for. Nevertheless, estimates of prediction accuracy calculated using data from a highly replicated, independent trial of only accessions with previously determined acceptable performance (i.e., minimal shattering and lodging) also gave an optimistic outcome for using predictions to assist in the selection of superior accessions.

Based on a comparison of predictions and observed field performance in each trial, a soybean breeder could select the top 10% of accessions based on genomic prediction of yield and expect 88% of the selected accessions to be better than average for yield. This example demonstrated that genomic predictions can be used to enrich field trials of accessions with accessions that perform better than a randomly selected set. Looking at the extremes, we found that the top 10% for each trait rarely contained accessions that performed in the bottom 10% according to actual trial data, indicating that using predictions very effectively eliminates the accessions that hold little promise, ultimately saving field resources to evaluate more of those that do hold promise.

Compiling historical data on accession evaluations conducted across four states going back to 1963 provided us a very large training dataset consisting of over 9,000 accessions. Soybeans adapted to different latitudes belong to different MGs. The trials used as a source of data ranged from trials conducted on early MGs in Minnesota to late MGs conducted in MS. We explored the optimal use of such a large and diverse training set for calibrating genomic prediction models. Our results can be summarized by two basic findings. First of all, the population and target environment being predicted should be well represented in the training set. The poorest predictive abilities were observed when we attempted to predict between states or between genetic clusters. Secondly, genomic prediction training sets appear to be very forgiving to the presence of data from diverse geographical locations and genetic clusters. It was surprising to observe that adding data from very different geographical locations had no effect on predictive ability. For example, the prediction of performance in MN environments was not affected by the presence of training data collected in MS on MG VII - IX accessions. This may partially be an artifact stemming from the tendency of accessions from similar MGs to genetically cluster, and the partitioning of MGs across the states used for evaluation. In BLUP, information from closer relatives is weighted more heavily, while less weight is given to information from distant relatives (de Los Campos et al., 2013), meaning data from MS was probably weighted less heavily in the prediction of early MG accession performance in MN.

Building a training set by adding accessions from different and diverse genetic clusters did not improve nor harm predictive ability when the goal was to predict accession performance within a single cluster. One exception included the prediction of yield corrected for shattering and lodging across diverse clusters. Our general results are not consistent with results from barley that suggested that the addition of unrelated individuals to a training set can potentially reduce predictive ability (Lorenz and Smith 2015), but they are consistent with results in maize where training sets were formed by combining data across heterotic groups (Technow *et al.* 2013). The underlying reasons for the neutral effect of adding genetically distant individuals to the soybean accession training sets could relate to the flow of information from historical LD and pedigree relationships to prediction accuracy (Habier *et al.* 2013). In the barley case (Lorenz and Smith 2015), where there is substantive family structure and a high degree of relatedness among lines from the same breeding program, it is likely that pedigree relationships, captured by G, are the predominant source of accuracy. The addition of less related individuals can reduce the accuracy provided by this source of information (Habier *et al.* 2013). In the case of the soybean germplasm collection, where many individuals do not share close pedigree relationships and where common ancestors likely go back many generations, the predominant source of accuracy is likely historical LD. The large training populations and high marker densities may have allowed the capturing of this information (Habier *et al.* 2013; Hickey *et al.* 2014), offsetting any possible detrimental effect on the genetic relationships source of information.

In conclusion, this study demonstrates that historical data collected as part of plant germplasm collection characterizations can be used to develop predictive models to help breeders select accessions for introgressing useful genetic variation. We found that in the case of the soybean germplasm collection, these models are robust to the inclusion of diverse sources of data, but training sets should include data from populations and environments representative of the target populations and environments. This data has already been collected and made freely available, and therefore nothing is preventing the use of these models for enhancing utilization of this genetic resource. Genomic predictions might also be used to develop trait-specific “core collections” that could be used for deeper phenotyping for detailed studies on physiological mechanisms and high-resolution QTL mapping. It is anticipated that the genomics revolution will create similar data resources for germplasm collections of other agriculturally important species and that genomic prediction will serve as a key tool for making practical use of the genomic data.

---

**Supplementary Figure 1.**
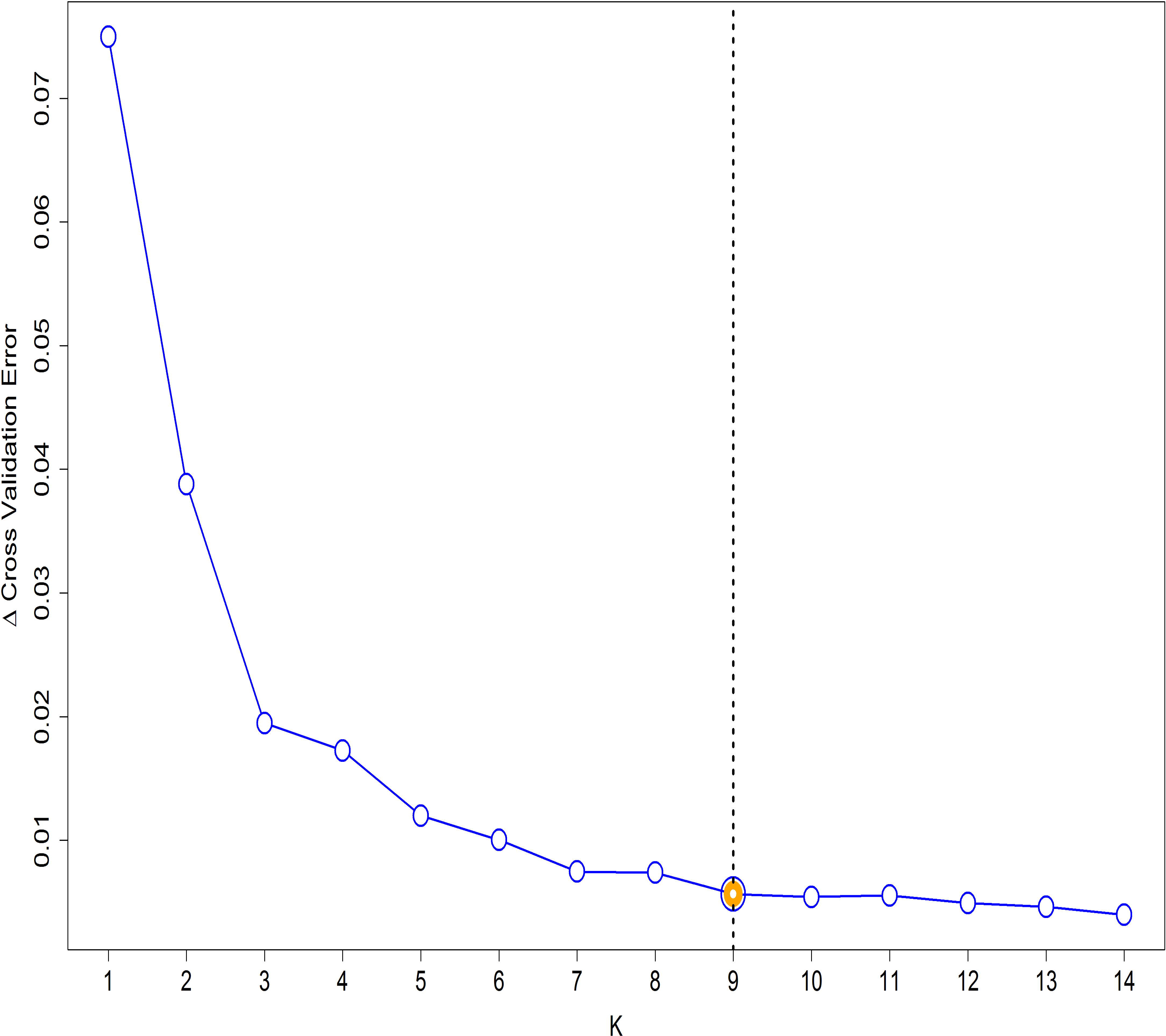
Exploration of the optimal number of genetic subpopulations (K) within the set of soybean accessions included in this study. A difference in cross-validation error between levels of K was used as a criteria. A plateau in Δ cross-validation error at K=9 was used to infer K.

## Supplementary Table Captions

Table S1. Predictive abilities from the *One/Group*, *Group/All,* and *One/All* cross validation schemes for oil, protein, and yield in each trial using the G-BLUP model. The grouping factor was Trial.

Table S2. Predictive ability for seed yield using the *Group/All, One/All,* and *One/Group* cross validation schemes. The grouping factor is Trial. Tabled values are predictive abilities when phenotypes are not adjusted for early shattering or lodging (None), adjusted for lodging (L), adjusted for early shattering (S), or adjusted for both (B). Only trials for which early shattering and lodging data were available were included in this analysis.

Table S3. Predictions for *Group/Group* cross validation scheme with data grouped by trial. The cells are shaded according to the value of the correlation coefficient with red shades indicating higher correlations and blue shades indicating lower correlations relative to the average correlation.

Table S4. Number and percentage of accessions belonging to each cluster separated by maturity group.

Table S5. Raw phenotypes, corrected phenotyped, and predictions of phenotyped accessions comprising this study.

Table S6. Genomic predictions (G-BLUP) of non-phenotyped accessions contained within the USDA Soybean Germplasm Collection.

